# UHRF1 is critical for tumor-promoting inflammation and tumorigenesis in retinoblastoma

**DOI:** 10.1101/2025.03.02.641083

**Authors:** Yijun Gu, Marina Suarez-Pizarro, Claire C. Chen, Jie Wu, Loredana Zocchi, Susie Suh, Parker J. Smith, Sylvía Cruz, Roberto Tinoco, Claudia A. Benavente

**Affiliations:** Department of Pharmaceutical Sciences, University of California, Irvine, CA 92697, USA; Department of Developmental and Cell Biology, University of California, Irvine, CA 92697, USA; Department of Biological Chemistry, University of California, Irvine, CA 92697, USA; Chao Family Comprehensive Cancer Center, University of California, Irvine, CA 92697, USA; Gavin Herbert Eye Institute, Department of Ophthalmology, University of California, Irvine, CA 92697, USA; School of Biological Sciences, University of California, Irvine, CA 92697, USA; Department of Molecular Biology and Biochemistry, University of California, Irvine, CA 92697, USA

## Abstract

Retinoblastoma, the most common pediatric intraocular malignancy, arises from RB1 inactivation, leading to uncontrolled proliferation of retinal progenitor cells. Epigenetic dysregulation a key driver of retinoblastoma progression, yet the underlying mechanisms are poorly understood. UHRF1, a regulator of DNA methylation and chromatin remodeling, has been implicated in oncogenesis but its role in retinoblastoma has not been fully characterized. To investigate Uhrf1’s role in tumor initiation and progression, we generated a genetically engineered mouse model of retinoblastoma with conditional *Uhrf1* knockout. Remarkably, Uhrf1 loss completely blocked tumor formation, despite persistent early oncogenic events. Transcriptomic and epigenomic profiling revealed that Uhrf1 is essential for tumor progression, facilitating oncogenic transcription, chromatin accessibility, and aberrant DNA methylation. Beyond its epigenetic role, we found Uhrf1 modulates the tumor immune microenvironment, promoting chemokine secretion and microglial infiltration, suggesting that Uhrf1 promotes immune cell recruitment. Mechanistically, Uhrf1 enhances chemokine expression through NF-κB signaling, establishing a novel connection between epigenetic regulation and tumor-associated immune responses. These findings establish Uhrf1 as a critical driver of retinoblastoma progression, essential for tumor maintenance but not initiation. By sustaining oncogenic transcriptional programs and shaping a pro-tumor immune microenvironment, Uhrf1 emerges as a promising therapeutic target for inhibiting tumor growth and immune evasion.

## Introduction

The retinoblastoma tumor suppressor (RB1) is a master cell cycle regulator. Mutations or functional inactivation of RB1 occur in the majority of human cancers. Germline mutations in the *RB1* gene result in retinoblastoma, a rare form of childhood cancer, and increase the risk of osteosarcomas and other cancers (Friend et al. 1986). Retinoblastoma tumors typically have stable genomes, with tumor progression primarily driven by epigenetic dysregulation of various cancer pathways (Zhang et al. 2012). However, the key factors driving epigenome changes following *RB1* inactivation, especially in the developing retina, remain unclear. Recent studies identified Hells (helicase, lymphoid specific) as a significant contributor to retinoblastoma tumor initiation and progression (Benavente et al. 2014; Zocchi et al. 2020a). However, Hells functions as a transcriptional co-activator of E2F3 to promote tumorigenesis rather than driving the epigenome restructuring required for retinoblastoma formation (Zocchi et al. 2020a, 2020b).

Retinogenesis involves several critical stages, from the proliferation of retinal progenitor cells (RPCs) to their differentiation and maturation into various retinal neurons and Müller glial cells. In the mouse retina, a common population of RPCs emerges between embryonic day 11 (E11) and postnatal day 10 (P10). These cells undergo unidirectional differentiation in a conserved temporal and spatial order (Figure 1A) (Cepko 2014). This process is regulated by complex molecular mechanisms, including transcriptional and post-transcriptional regulation. While transcription factors governing specific neurogenic fates have been extensively studied, much less is known about the epigenetic events involved. Studies of mammalian neurogenesis, including retinal development, reveal gradual changes in the chromatin states of RPCs (Lister et al. 2013; Ziller et al. 2015; Mo et al. 2015, 2016; Aldiri et al. 2017). Alterations in epigenetic markers during retinal development have been linked to various eye diseases, including retinoblastoma (Zhang et al. 2012; Benavente et al. 2014; Zocchi et al. 2020a; Aldiri et al. 2017; Cvekl and Mitton 2010; Pennington and DeAngelis 2015). Understanding the chromatin factors that regulate retinal stem cell maintenance and neurogenesis is vital for understanding normal retinal development, identifying mechanisms underlying retinal disorders, and discover potential therapeutic targets.

**Figure 1.**
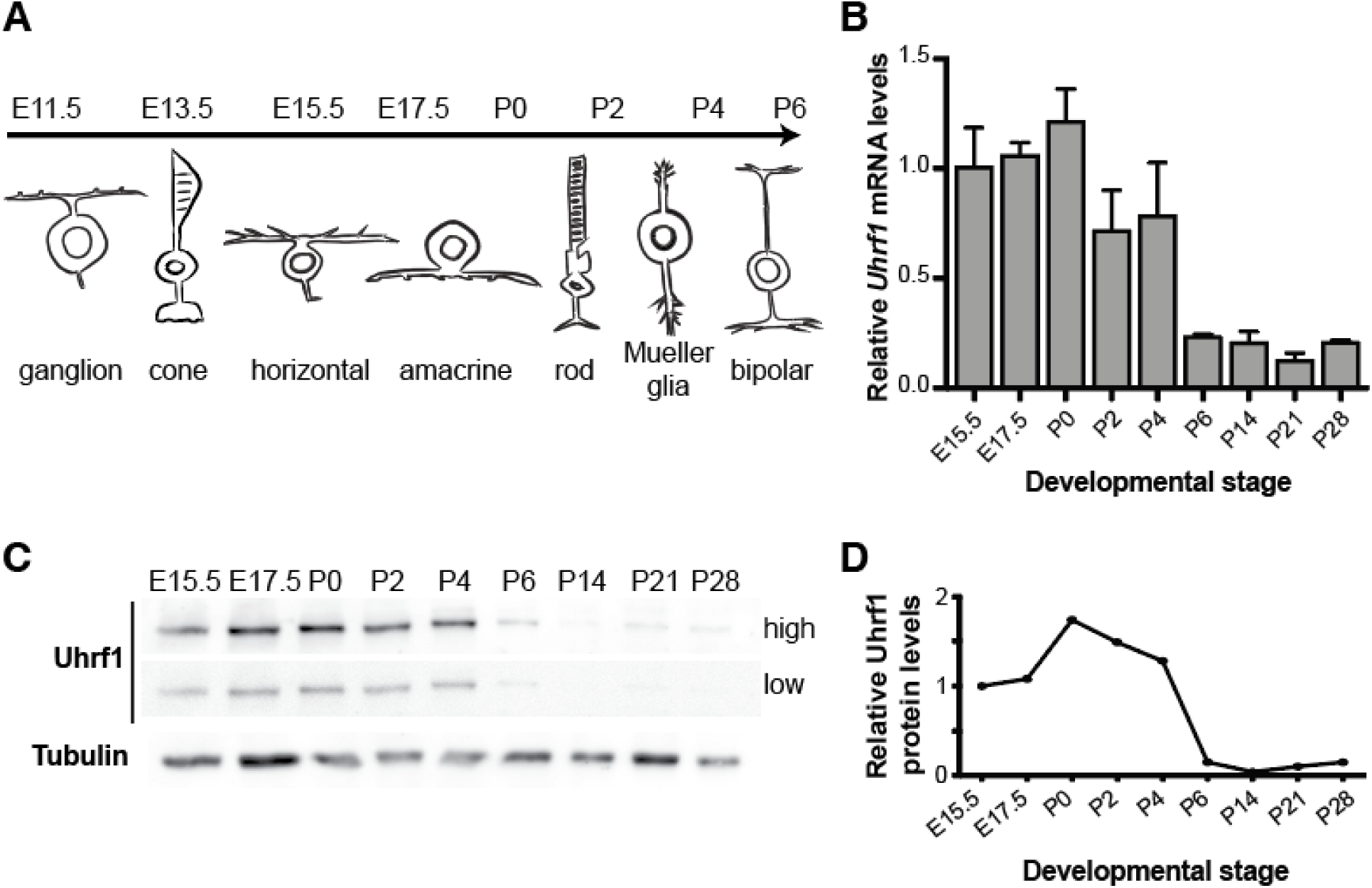
Uhrf1 is repressed during the late stages of retinal development. (A) Illustration of the developmental stages and specification timing of the 7 major retinal cell types: retinal ganglion cells, amacrine, horizontal, bipolar, Müller, cone, and rod photoreceptors. (B) Time course of *Uhrf1* mRNA expression in the developing retina in wild-type mice. Levels of *Uhrf1* mRNA were measured by RT-qPCR and normalized to the levels at E15.5 taken as 1 (n=3). Mean ± SD. (C) Representative Western blot analysis of UHRF1 protein levels with high and low exposure. Tubulin was used as loading control. (D) Quantification of the relative UHRF1 protein levels on Western blots, were E15.5 was taken as 1 (n=3). Mean ± SD.

UHRF1 (ubiquitin-like with PHD and RING finger domains 1) is a critical mediator of DNA methylation, histone modification, and chromatin remodeling. It predominantly interacts with DNA methylatransferase 1 (DNMT1), playing a central role in maintaining DNA methylation in dividing cells. UHRF1 also contains a RING finger domain with E3 ligase activity, facilitating to histone and protein ubiquitination (Citterio et al. 2004; Nishiyama et al. 2013; Liu et al. 2013; Qin et al. 2015). Additionally, UHRF1 interacts with histone deacetylase 1 (HDAC1) and histone methyltransferases (G9a and Suv39H1), to establish and reorganize heterochromatin (Kim et al. 2009). In non-cancerous cells, UHRF1 is expressed during proliferation and downregulated upon differentiation, with expression fluctuating throughout the cell cycle (Sharif et al. 2007; Sen et al. 2010; Tien et al. 2011). In contrast, UHRF1 expression is sustained throughout all cell cycle phases in cancer cells and plays a pivotal role in cancer formation and progression (Tien et al. 2011; Alhosin et al. 2011; Hervouet et al. 2010; Babbio et al. 2012; Pacaud et al. 2014; Wu et al. 2022b; Kim and Benavente 2024). Furthermore, the *UHRF1* gene is transcriptionally repressed by RB1 and is implicated in the epigenetic changes associated with retinoblastoma, making it critical for tumor survival (Benavente et al. 2014; Guzman et al. 2020). However, whether UHRF1 is necessary for retinoblastoma development or if it is a critical mediator of the epigenome changes that follow *RB1* inactivation in the retina, remains unexplored.

In this study, we investigate whether UHRF1 is required for tumor initiation, progression, or both in a genetic mouse model of retinoblastoma. Using genetically engineered mouse models (GEMMs) of retinoblastoma, we show that loss of Uhrf1 prevents tumor formation, despite the persistence of early oncogenic events. Through transcriptomic and epigenomic analyses, we demonstrate that Uhrf1 overexpression drives tumor progression by sustaining oncogenic transcriptional programs and altering chromatin accessibility and DNA methylation. Additionally, we uncover a role for Uhrf11 in shaping the tumor immune microenvironment, promoting chemokine secretion and immune cell recruitment via NF-κB signaling. Our findings reveal Uhrf1 as a key epigenetic regulator in retinoblastoma, driving both the tumor-supportive transcriptional landscape and immune infiltration.

## Results

### Uhrf1 is highly expressed during the proliferative stages of retinal development and downregulated upon terminal differentiation

To investigate the expression pattern of Uhrf1 in the developing retina, we performed real-time reverse transcriptase PCR (RT-qPCR) and Western blot analyses on retinal tissue at different developmental stages. The analysis revealed that *Uhrf1* mRNA is robustly expressed starting at embryonic day 15.5 (E15.5), peaks around postnatal day 0 (P0), and gradually declines until reaching minimal levels by postnatal day 6 (P6), where it remains low thereafter (Figure 1B). The mRNA levels closely correlate with Uhrf1 protein expression. High levels of UHRF1 protein are observed during the proliferative stages of retinal development (E15.5-P2). A steep decline in protein expression occurs after this phase, falling below the detection limit following the completion of retinal cell-fate specification at P6 (Figures 1C-D).

### Uhrf1 is dispensable for normal retinal development

To elucidate the role of Uhrf1 during retinal development, we generated genetically engineered mice with the *Uhrf1* gene knockout (Figure 2A) in early retinal progenitors using the *Chx10-Cre* allele (*Chx10-Cre; Uhrf1^lox/lox^*; referred hereafter as *Uhrf1* cKO). Some of these mice also carried the *Z/EG* transgene as a Cre-recombination reporter. Western Blot analysis of P0 retinae confirmed a significant reduction in UHRF1 protein levels in *Uhrf1* KO mice compared to *Cre*- negative littermate controls (Figure 2B). Since *Chx10-Cre* is expressed stochastically, complete loss of protein expression was not expected in the entire retina.

**Figure 2.**
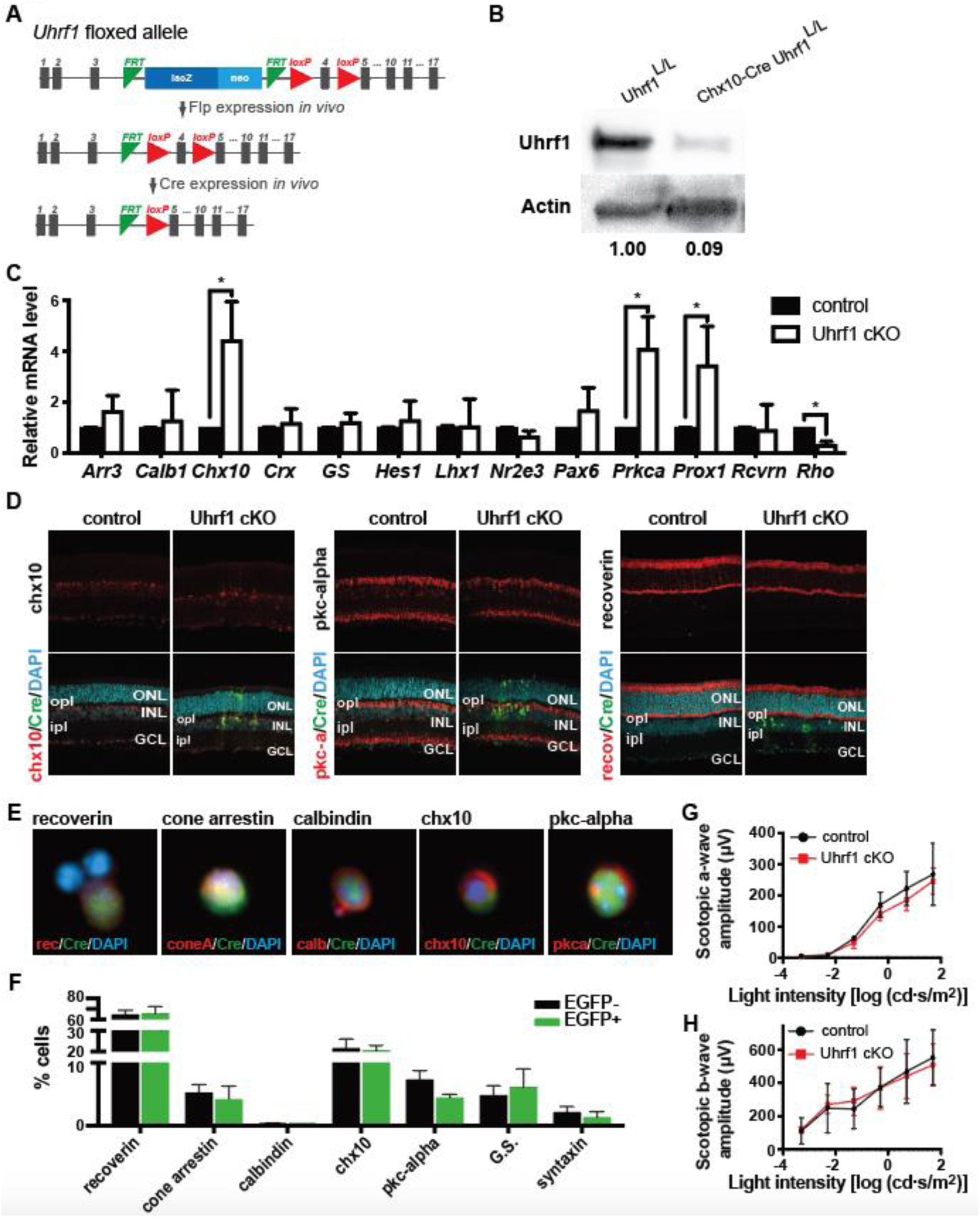
*Uhrf1* is nonessential for normal retinal development. (A) Schematic diagram of conditional excision of the floxed *Uhrf1* allele. (B) Western blot analysis of Uhrf1 expression in *Uhrf1* cKO (Chx10-*Cre Uhrf1*^lox/lox^) and littermate control (*Uhrf1*^lox/lox^) shows effective reduction of Uhrf1 protein upon Cre recombination of floxed *Uhrf1* alleles. Actin was used as loading control. (C) RT-qPCR analysis of retinal cell marker genes. A significant increase in the mRNA level of *Chx10* (bipolar), *Prkca* (bipolar)*, Prox1* (amacrine/horizontal) and a significant decrease in *rhodopsin* (rods) transcription were observed in *Uhrf1* cKO EGFP+ cells compared to the control littermates. All data are normalized to control littermates (n=4). *p<0.05 by unpaired two-tailed t test. (D-E) Representative images of P21 retina cross-sections (D) and dissociated cells (E) from *Uhrf1* cKO *Z/EG* mice immunostained with recoverin (photoreceptors), cone-arrestin (cone photoreceptors), calbindin (horizontal and a subset of amacrine cells), chx10 (bipolar), and pkc-alpha (bipolar) antibodies (red). Retinae were double immunostained with anti-GFP to capture areas of *Chx10-Cre*-mediated GFP expression. Nuclei were counterstained with DAPI (blue). ONL, outer nuclear layer; INL, inner nuclear layer; GCL ganglion cell layer; ipl, inner plexiform layer; opl, outer plexiform layer. (F) Quantification of the proportion of immunoreactive cells for each cellular marker antibody shown in E and supplemental figure 3 was determined for EGFP+ and EGFP-cells from *Uhrf1* cKO *Z/EG* mice from independent litters (n=3). Each bar represents the mean ± SD of 500 cells scored from each retina. (G-H) ERGs were recorded from 5-week-old *Uhrf1* cKO (red line) and littermate control (black line). a-wave amplitude (G) and b-wave amplitude (H) were recorded at various light intensities. All measurements are mean ± SD (n=4).

We used RT-qPCR to assess the expression of retinal cell markers for the seven major retinal cell types in P21 retinae. There were no significant differences in the expression of *Arr3*, *Calb1*, *Chx10*, *Crx*, *GS*, *Hes1*, *Lhx1*, *Nr2e3*, *Pax6*, *Prkca*, *Prox1*, *Rcvrn*, and *Rho* between *Uhrf1* cKO and *Cre*-negative controls (Figure 2C). This suggests that Uhrf1is not required for normal retinal development.

Retinal cross-sections were examined using immunohistochemistry (Figure 2D and Supplemental Figure 2) and dissociated retinae using immunocytochemistry (Figure 2E and Supplemental Figure 3). Immunostaining with antibodies specific to major retinal cell types revealed no significant abnormalities in cell birth or cell-fate specification (Supplemental Figure 2). The outer nuclear layer (ONL), inner nuclear layer (INL) and ganglion cell layer (GCL) displayed normal structure, uniform thickness, and correct cellular ratios (Figure 2D and Supplemental Figure 2). The only observed abnormality was in bipolar cells (Chx10-positive cells). Bipolar cell processes in *Uhrf1* cKO retinae extended through the outer plexiform layer (OPL) into the ONL, whereas in the *Cre*-negative regions, bipolar cell bodies remained confined to the INL (Figure 2D).

Immunocytochemical analysis of dissociated retinae confirmed that all retinal cell types were present in the correct ratios, as double-stained cells for each retinal cell-specific marker and GFP reporter were detected (Figure 2E-F and Supplemental Figure 3). RT-qPCR analysis of EGFP+ retinal cells sorted by flow cytometry further corroborated the absence of significant differences in retinal marker expression (Supplemental Figure 1).

Electroretinography (ERG) was conducted on dark-adapted *Uhrf1* cKO mice with retinae exhibiting at least 50% EGFP+ penetrance to evaluate retinal function. ERG measures the electrical activity of retinal cells in response to light stimulation. The a-wave, which reflects photoreceptor (cone and rod) function, showed no significant differences in amplitude (Figure 2G). The b-wave, representing inner retinal function mediated predominantly by Müller and bipolar cells, also displayed no significant changes in amplitude (Figure 2H).

These findings demonstrate that *Uhrf1* is dispensable for normal retinal development. Despite the minor structural abnormality in bipolar cell processes, all retinal cell types are born, terminally differentiate, and function within normal parameters in the absence of Uhrf1.

### Uhrf1 is transcriptionally repressed by the Rb family

Previous studies have demonstrated that *UHRF1* is regulated through the RB/E2F pathway, with loss of *RB1* leading to UHRF1 upregulation and overexpression. This process may contribute to epigenetic changes driving tumor progression (Benavente et al. 2014; Wu et al. 2022b). We confirmed Uhrf1 overexpression in the adult retina (P21) of *Chx10-Cre Rb1^lox/lox^ Rbl1^-/-^* (*Rb1/Rbl1* DKO) mice (Figure 3A) as well as in retinoblastoma tumors (Figure 3B). Chromatin immunoprecipitation (ChIP) analysis using P0 mouse retinae and the human retinoblastoma cell line (Weri) confirmed E2F1 enrichment at consensus-binding motifs within the *Uhrf1* promoter (Figure 3C).

**Figure 3.**
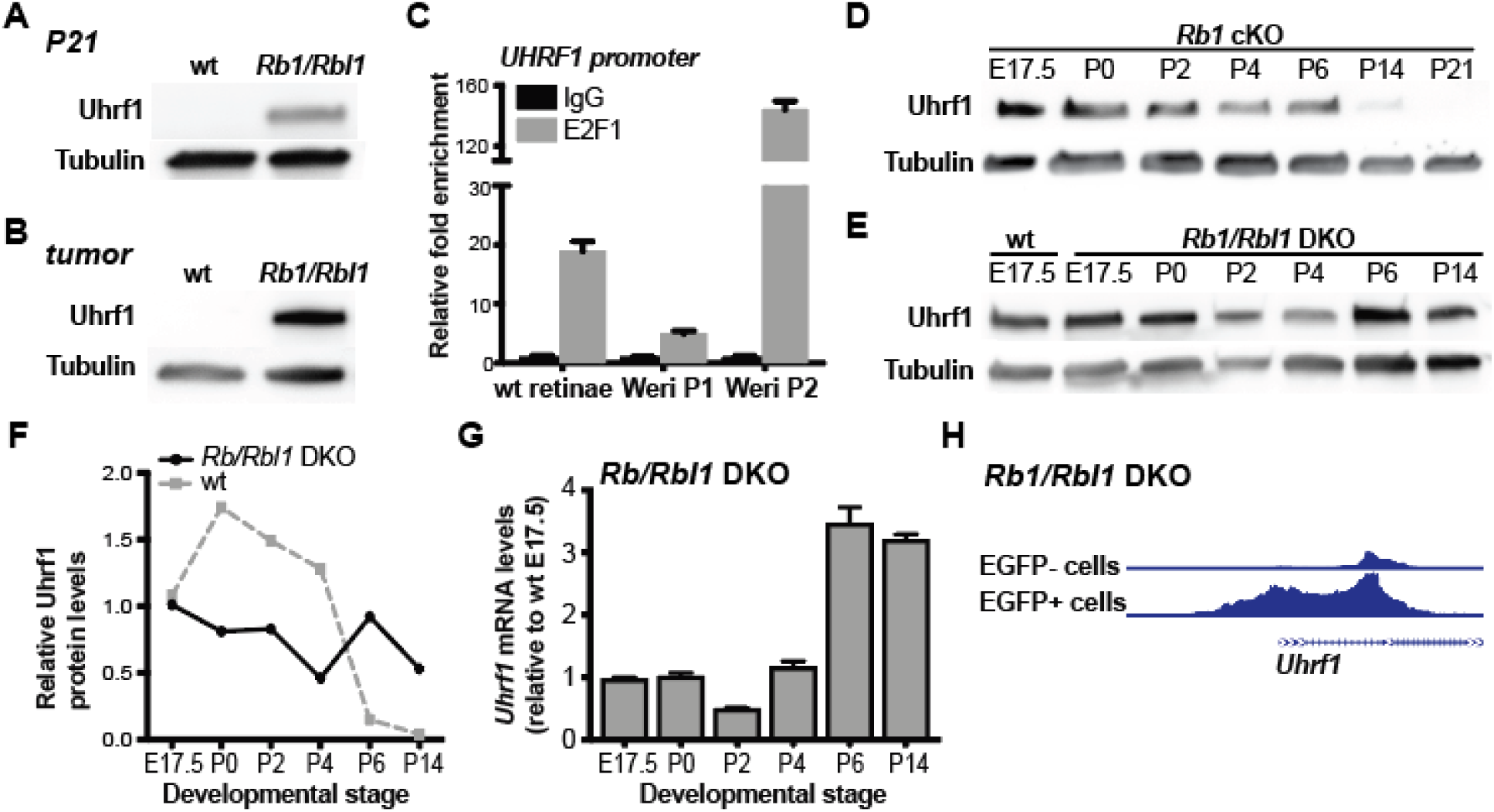
Uhrf1 is transcriptionally repressed by RB1 and RBL1. (A-B) Uhrf1 protein level in P21 retinae and retinoblastoma tumor samples were compared to littermate controls (wt). Tubulin was used as loading control. (C) Chromatin immunoprecipitation (ChIP) assay in P0 mouse retinae and Weri retinoblastoma human cell line reveals enrichment of E2f1 within the *Uhrf1* promoter. (D-E) Western blot analysis of Uhrf1 protein level in different retinae development stages in *Rb1* cKO mice and *Rb1/Rbl1* cDKO. Tubulin was used as loading control. (F) Quantification of Uhrf1 protein expression in *Rb/Rbl1* cDKO. (G) RT-qPCR analysis of *Uhrf1* mRNA level in *Rb1/Rbl1* cDKO. *Uhrf1* mRNA level in E17.5 wt mouse was set as 1. (H) Representative sequencing tracks for the *Uhrf1* locus show increased peaks at the promoter in P21 *Rb1/Rbl1* DKO mouse retinal cells (EGFP+) compared to control retinal cells (EGFP-). The ATAC-Seq data have been normalized to take sequencing depth into account. All measurements are mean ± SD (n=3).

To assess whether Uhrf1 overexpression occurs immediately after RB family loss or specific retinal cell types, we analyzed Uhrf1 expression in *Chx10-Cre Rb1*^lox/lox^ (*Rb1* cKO) and *Rb1/Rbl1* DKO retinae. In *Rb1* cKO retinae, which do not develop retinoblastoma due to compensation by *Rbl1*, Uhrf1 mRNA and protein expression followed a similar pattern to wild-type retinae. Expression peaked during embryonic stages (E17.5-P0), was repressed after postnatal day 2 (P2) and showed significant reduction by P6 (Figure 3D). In contrast, P21 retinae from *Rb1/Rbl1* DKO mice exhibited increased Uhrf1 protein levels compared to littermate controls (wt; Figure 3A). Further analysis of *Rb1/Rbl1* DKO retinae at various stages showed the absence of UHRF1 repression at both protein (Figure 3E-F) and transcriptional levels (Figure 3G), compared to wild-type controls.

ATAC-seq analysis of chromatin accessibility in P21 retinae revealed a significant increase in reads at the *Uhrf1* promoter and the first two exons in EGFP-positive (Rb1/Rbl1 null) compared to EGFP-negative (wt-like) flow-sorted *Rb1/Rbl1* DKO retinal cells (Figure 3H).

### Loss of Uhrf1 prevents tumor progression in a genetic mouse model of retinoblastoma

To determine whether Uhrf1 overexpression is essential for retinoblastoma development, we generated *Chx10-Cre Rb1^lox/lox^ Rbl1^-/-^ Uhrf1 ^lox/lox^* triple knockout (*Rb1/Rbl1/Uhrf1* TKO) mice and compared them to *Rb1/Rbl1* DKO littermate controls. *Rb1/Rbl1/Uhrf1* TKO and *Rb1/Rbl1* DKO mice were monitored from birth for one year to assess tumor development. Remarkably, abrogation of *Uhrf1* resulted in 100% tumor-free survival in *Rb1/Rbl1/Uhrf1* TKO, whereas 65.4% of *Rb/Rbl1* DKO mice developed significant tumors, as previously reported (Zhang et al. 2004b; Benavente et al. 2014; Zocchi et al. 2020a; Benavente et al. 2013) (Figure 4A).

**Figure 4.**
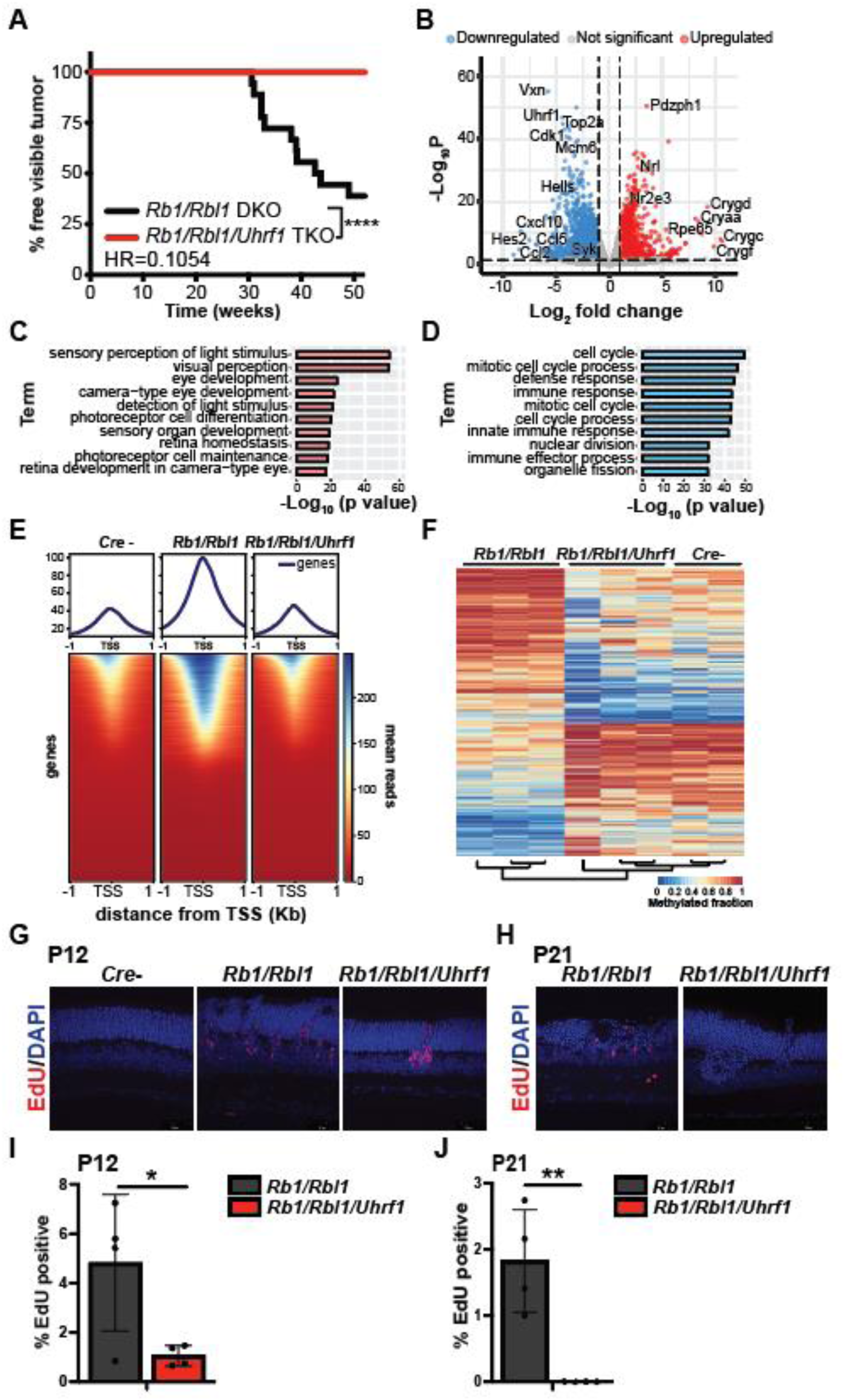
Loss of *Uhrf1* prevents retinoblastoma formation. (A) Kaplan-Meier curves showing the percentage of mice free of visible retinoblastoma tumors in *Rb1/Rbl1* DKO (n=52) and *Rb1/Rbl1/Uhrf1* (n=41) mice. (B) Volcano plot of differentially expressed genes (DEGs) in *Rb1/Rbl1/Uhrf1* TKO compared to *Rb1/Rbl1* DKO P21 mouse retinae. (C-D) p-value ranking bar graph representing the ten most significant gene ontology (GO) biological processes for (C) upregulated genes and (D) downregulated genes in *Rb1/Rbl1/Uhrf1* TKO compared with *Rb1/Rbl1* DKO P21 retinae. (E) Genome-wide heatmap plot of chromatin accessibility peaks (ATAC-seq) from flow sorted EGFP-negative (*Cre*-negative) *Rb1/Rbl1/Uhrf1* and EGFP-positive *Rb1/Rbl1* DKO and *Rb1/Rbl1/Uhrf1* TKO P21 retina grouped by mean reads and by distance from the transcription starting site (TSS). Each row represents a gene ordered in descending accessibility mean reads. (F) Heat map of differentially methylated genes in *Cre*-positive *Rb1/Rbl1/Uhrf1*, *Rb1/Rbl1* DKO and *Cre*-negative *Rb1/Rbl1/Uhrf1* TKO P21 retina. (G-H) Representative images of (G) P12 retina and (H) P21 retina cross-sections from *Rb1/Rbl1* DKO and *Rb1/Rbl1/Uhrf1* TKO mice labeled with EdU (proliferating cells; red). Nuclei were counterstained with DAPI (blue). (I-J) Quantification of the proportion of EdU positive cells in (I) P12 and (J) P21 *Rb/Rbl1* cDKO and dissociated retina Each bar represents the mean ± SD of 500 cells scored from each retina. (n=3). *p<0.05, **p<0.01 by unpaired two-tailed t test.

To explore the molecular changes underlying Uhrf1-driven tumor progression, we performed RNA-seq analysis on P21 retinae from *Rb1/Rbl1/Uhrf1* TKO and *Rb1/Rbl1* DKO mice. This analysis identified a total of 1444 differentially expressed genes (DEGs), including 913 upregulated and 531 downregulated genes in *Rb1/Rbl1/Uhrf1* TKO retinae compared to *Rb1/Rbl1* DKO retinae (Figure 4B). When DEGs were grouped into upregulated and downregulated, genes related to visual system-related signaling and functions, and photoreceptor cells contained the highest enrichment among upregulated genes (Figure 4C and Supplemental Figure 4). Conversely, genes related to cell cycle replication and immune response pathways and horizontal cells were enriched among downregulated genes (Figure 4D and Supplemental Figure 4).

Given Uhrf1’s established role in maintaining DNA methylation and chromatin structure, we further examined the impact of Uhrf1 loss on chromatin accessibility and DNA methylation. ATAC-seq analysis of flow-sorted EGFP+ P21 retinal cells from *Rb1/Rbl1/Uhrf1* TKO and *Rb1/Rbl1* DKO mice carrying the Z/EG reporter gene revealed distinct chromatin accessibility patterns. As previously observed, *Rb1/Rbl1* DKO retinae exhibited increased chromatin accessibility, consistent with tumor-prone epigenomes (Figure 4E). Interestingly, *Rb1/Rbl1/Uhrf1* TKO retinae displayed chromatin accessibility patterns resembling those of *Cre-* negative controls, suggesting a restoration of normal chromatin structure (Figure 4E). To assess the effect of Uhrf1 loss on DNA methylation, we performed reduced representation bisulfite sequencing (RRBS) to analyze differentially methylated regions across the genome. This analysis revealed widespread changes in DNA methylation in *Rb1/Rbl1* DKO retinae. Importantly, these epigenetic changes were reversed in *Rb1/Rbl1/Uhrf1* TKO retinae, whose methylation profiles closely resembled those of *Cre*-negative controls (Figure 4F).

Since the loss of Uhrf1 restores normal transcriptomic and epigenomic profiles, effectively reversing tumorigenesis, a critical question remains: Is Uhrf1 required for tumor initiation by creating a permissive epigenetic landscape for transformation, or does it primarily function later to sustain tumor progression through oncogenic transcriptional programs? To investigate this, we examined cellular proliferation in the developing retina. In the developing mouse retina, RPCs normally cease proliferation by P10 (Blanks and Bok 1977; Dyer and Cepko 2001). A hallmark of tumor initiation is the presence of ectopic proliferation beyond P10 (Chen et al. 2004; Zhang et al. 2004a). To address this, we performed EdU staining in P12 mice. As expected, no proliferating cells were detected in Cre-negative control retinae, whereas *Rb1/Rbl1* DKO retinae exhibited persistent proliferation (Figure 4G-I). Importantly, we also observed ectopic proliferation in *Rb1/Rbl1/Uhrf1* TKO retinae, though at a significantly lower frequency compared to *Rb1/Rbl1* DKO (Figure 4G-I). This suggests that Uhrf1 is not required for tumor initiation, as some degree of aberrant proliferation still occurs in its absence. However, while proliferating cells remain detectable at P21 in *Rb1/Rbl1* DKO retinae, no proliferating cells were observed in *Rb1/Rbl1/Uhrf1* TKO retinae at this stage. This suggests that while Uhrf1 is dispensable for tumor initiation, it is essential for sustaining long-term tumor growth.

Altogether, these findings indicate that Uhrf1 overexpression drives epigenetic changes that promote the transcription of pathways essential for tumor progression while inhibiting differentiation processes critical for retinal function. Loss of Uhrf1 restores normal epigenetic and transcriptomic profiles, effectively preventing tumor development in retinoblastoma-prone mice.

### Uhrf1 overexpression enhances recruitment of tumor-promoting microglia

Since genes involved in immune response pathways were highly enriched among downregulated genes (Figure 4D), we next investigated the nature of these immune responses. Deeper analysis of the transcriptomic data from P21 retinae of *Rb1/Rbl1/Uhrf1* TKO and *Rb1/Rbl1* DKO mice revealed a strong enrichment of cellular markers associated with microglia in *Rb1/Rbl1* DKO retinae (Figure 5A). Gene enrichment set analysis (GSEA) showed a reduced enrichment of fetal eye microglia gene markers in the absence of *Uhrf1*, suggesting that *Rb1/Rbl1* DKO retinae may experience higher microglial infiltration compared to Uhrf1-deficient retinae (Figure 5B).

**Figure 5.**
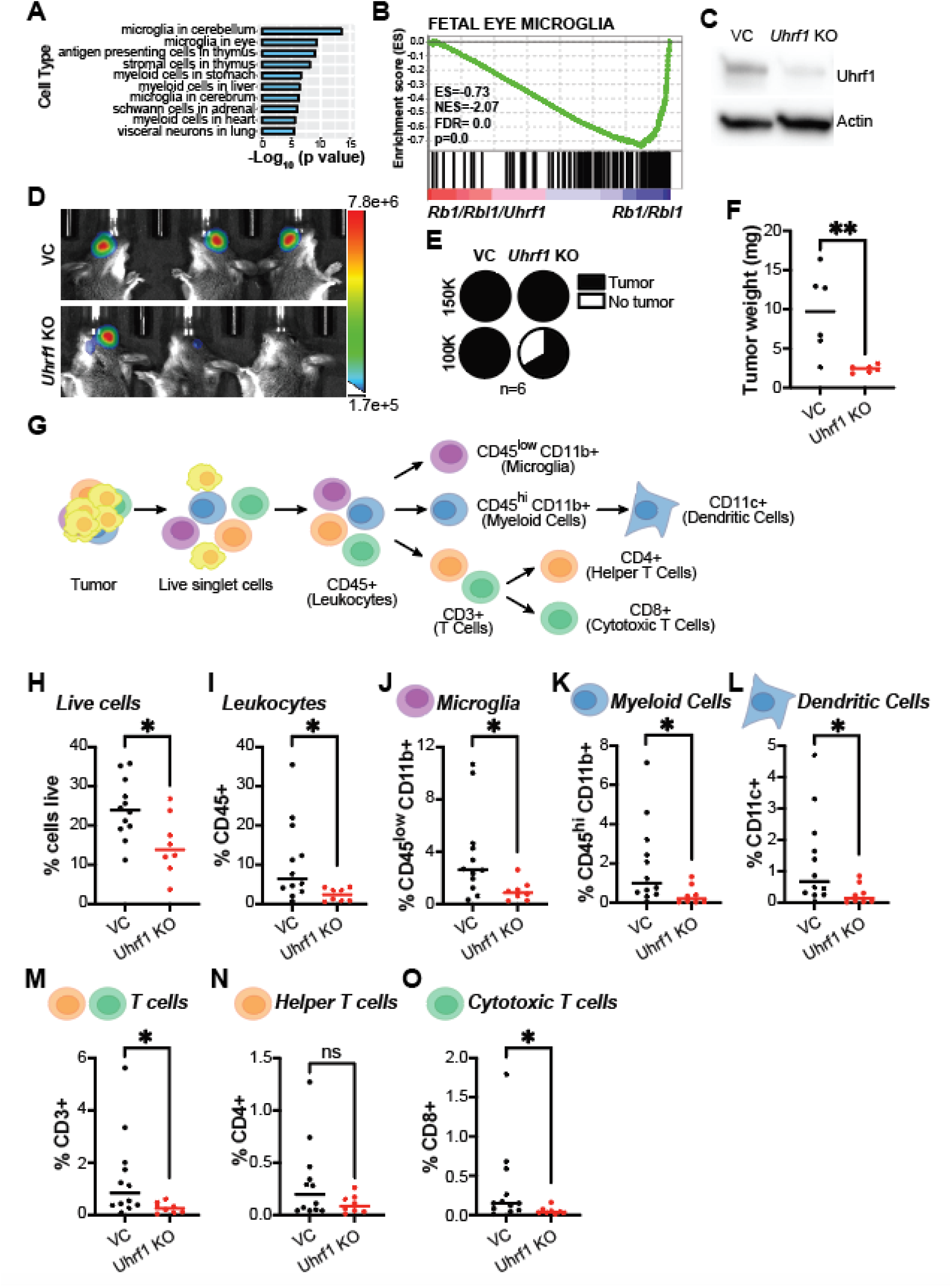
Uhrf1 overexpression enhances recruitment of tumor-promoting microglia. (A) p-value ranking bar graph representing the ten most significant cell types for downregulated genes in *Rb1/Rbl1/Uhrf1* TKO compared with *Rb1/Rbl1* DKO P21 retinae. (B) Gene Set Enrichment Analysis (GSEA) enrichment plot of the fetal eye microglia gene cluster from the RNA-seq data comparing *Rb1/Rbl1/Uhrf1* TKO to *Rb1/Rbl1* DKO P21 retinae. (C) Western blot analysis of Uhrf1 protein level in VC or *Uhrf1* KO UCI-Rb-1. Actin was used as loading control. (D) Representative bioluminescent images from mice orthotopically implanted with VC or *Uhrf1* KO UCI-Rb-1 cells at 2-weeks post sub-retinal injection. (E) Quantification of the percentage of mice presenting tumors at 2-weeks post implantation with either 150000 (150K) or 100000 (100K) cells in immune-competent mice. (F) Quantification of the final tumor weight at 2 weeks post implantation. (G) Schematic workflow of the immune flow analysis of retinoblastoma tumors derived from VC and *Uhrf1* KO UCI-Rb-1 cells after 2 weeks post-implantation. (H-O) Quantification of the percentage of (H) live cells, (I) leukocytes, (J) microglia, (K) myeloid cells, (L) dendritic cells, (M) T-cell, (N) helper T cells, and (O) cytotoxic T cells. Lines represent median ± SD. VC shown in black dots (n=12) and *Uhrf1* KO shown in red dots (n=8). For all graphs: ns=not significant, *p<0.05 by unpaired two-tailed t test.

Microglia, the resident immune cells of the retina, play a crucial role in coordinating immune responses against pathogens and damaged cells. Since photoreceptor degeneration is a key feature of the *Rb1*-null retina, we established the UCI-Rb-1 retinoblastoma cell line, derived from a *Rb1/Rbl1* DKO tumor, to minimize potential confounding effects. This model allowed us to directly assess whether Uhrf1 overexpression enhances tumor-immune interactions that support tumor progression. UCI-Rb-1 cells were genome edited using a CRISPR/Cas9 system with either a non-targeting guide RNA (gRNA) to be used as vector control (VC) or a *Uhrf1-* targeting gRNA (*Uhrf1* KO). Western blot analysis confirmed a significant reduction in Uhrf1 protein levels in *Uhrf1* KO clones compared to VC clones (Figure 5C). Both cell lines were transduced to express luciferase for in vivo tracking via bioluminescence imaging (BLI). VC and *Uhrf1* KO cells were orthotopically implanted into the sub-retinal space of *Cre*-negative *Rb1/Rbl1* DKO mice, followed by imaging at 24 hours and 2 weeks post-implantation (Figure 5D). At 24 hours, all mice displayed detectable tumor cell signals. However, by week 2, while all mice injected with 150000 cells showed evidence of tumors, only 66.7% (4/6) of mice injected with 100000 *Uhrf1* KO cells presented with tumors, compared to 100% of mice injected with VC cells (Figure 5D-E).

In follow-up studies, tumors were collected from luciferin-naïve mice two weeks post-implantation. At the time of collection, *Uhrf1* KO-derived tumors were significantly smaller (mean 2.40 ± 0.47 mg) compared to VC-derived tumors (mean 9.55 ± 5.24 mg; Figure 5F). To investigate changes in the tumor microenvironment, we analyzed the immune cell composition of VC- and *Uhrf1* KO-derived retinoblastomas (Figure 5G). VC-derived tumors had a significantly higher number of live cells than *Uhrf1*-KO (Figure 5H). Among these live cells, 10.71 ± 10.38 % were CD45-positive leukocytes in VC-derived tumors, whereas only 2.38 ± 1.65% were detected in *Uhrf1* KO tumors, indicating a significant reduction in immune cell infiltration in the absence of Uhrf1 (p=; Figure 5I). Microglia comprised the largest proportion of immune cells within the tumors. In VC-derived tumors, microglia accounted for 3.72 ± 3.36% of live singlet cells, or 38.62 ± 0.10% of CD45-positive cells. In contrast, *Uhrf1* KO-derived tumors had only 0.99 ± 0.77% microglia, representing 45.56 ± 0.19% of CD45-positive cells (p=0.0129; Figure 5J). Beyond microglia, *Uhrf1* KO tumors also showed robust decreases in other myeloid cell populations, including dendritic cells, and in cytotoxic T cells, but not helper T cells (Figure 5K-O).

### Uhrf1 activates NF-κB signaling to drive chemokine expression and immune cell recruitment

Given the dose-dependent nature of tumor implantation and the marked reduction in immune cell infiltration, particularly microglia, in *Uhrf1* KO-derived tumors, we next explored the mechanism by which Uhrf1 overexpression facilitates immune cell recruitment. Chemokines are key regulators of the tumor microenvironment, directing the migration of immune cells. To assess their involvement, we performed GSEA analysis on DEGs from *Rb1/Rbl1* DKO vs *Rb1/Rbl1/Uhrf1* TKO P21 retinae, focusing on the positive regulation of chemokine production gene set. The results revealed a reduced enrichment in the absence of *Uhrf1* (Figure 6A). To directly assess whether Uhrf1 modulates chemokine secretion, we analyzed the chemokine profiles of VC- and Uhrf1 KO cells. Our analysis revealed that retinoblastoma cells secrete CXCL1, CXCL10, CXCL12, CX3CL1, CCL2, CCL5, and CCL9, all of which were significantly reduced in *Uhrf1*-KO cells (Figure 6B-C). These chemokines are known to regulate immune cell recruitment, inflammation, and tumor progression. The overall reduction of these chemokines in *Uhrf1* KO cells suggests that Uhrf1 enhances chemokine secretion to facilitate microglial recruitment and immune cell infiltration, thereby fostering a tumor-supportive microenvironment.

**Figure 6.**
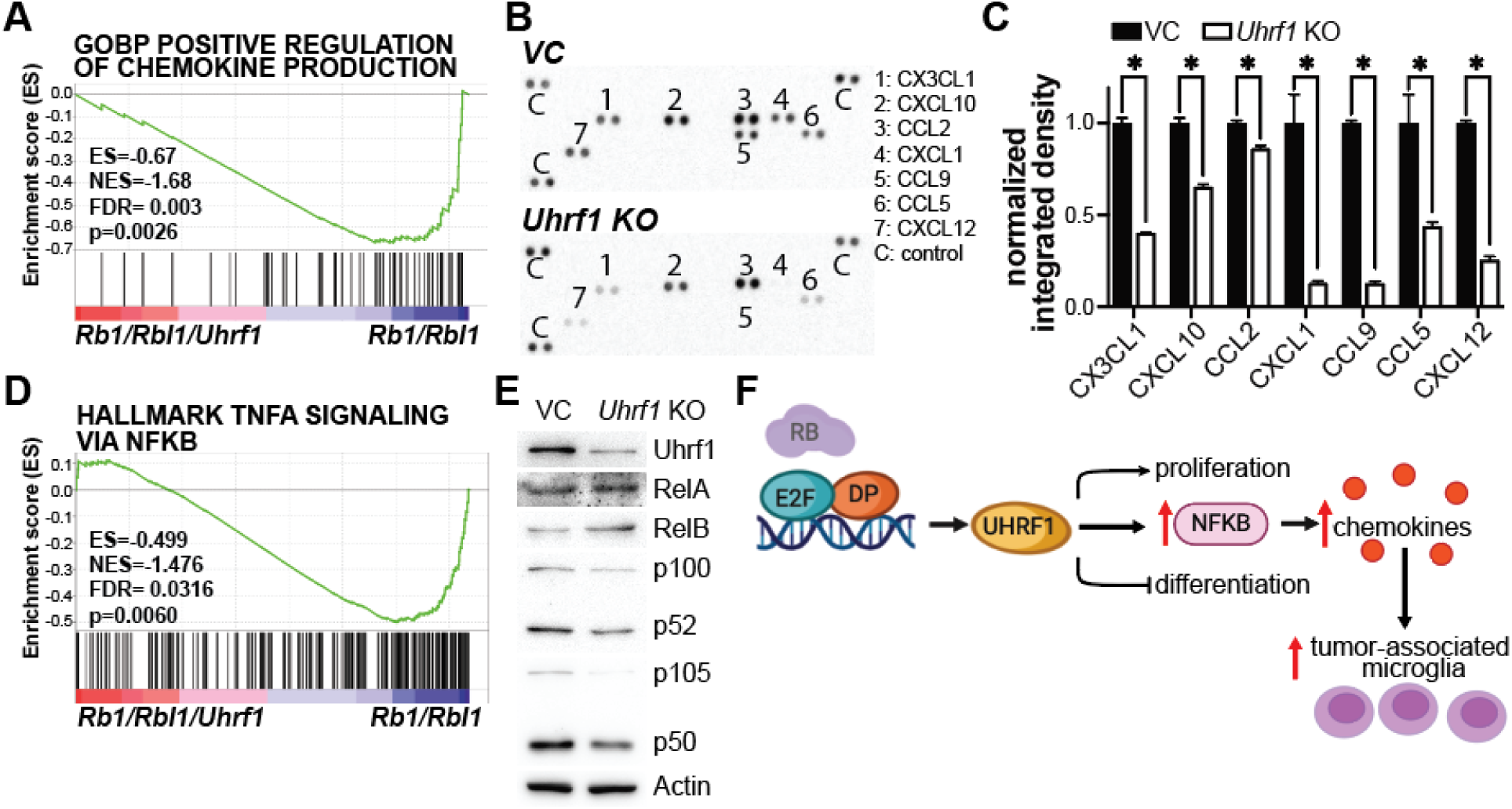
Uhrf1 loss decreases chemokine secretion and NF-κB expression. (A) Gene Set Enrichment Analysis (GSEA) enrichment plot of gene ontology for biological process for positive regulation of chemokine production from the RNA-seq data comparing *Rb1/Rbl1/Uhrf1* TKO to *Rb1/Rbl1* DKO P21 retinae. (B) Chemokine immunoassay analysis from conditioned media from VC and *Uhrf1* KO UCI-Rb-1 cells. (C) Quantification of the chemokines identified in B. (n=2 independent batches). mean ± SD *p<0.05 by unpaired two-tailed t test. (D) GSEA enrichment plot for the hallmark for TNF alpha signaling via NF-kB from the RNA-seq data comparing *Rb1/Rbl1/Uhrf1* TKO to *Rb1/Rbl1* DKO P21 retinae. (E) Western blot analysis of NF-kB family proteins level in VC and *Uhrf1* KO UCI-Rb-1. Actin was used as loading control. (F) Schematic summary of the role of Uhrf1 in retinoblastomagenesis.

Next, we sought to determine the underlying mechanism by which Uhrf1 regulates chemokine expression. Notably, all chemokines affected by UHRF1 loss are transcriptionally regulated by inflammatory signaling pathways, particularly NF-κB and STAT signaling. GSEA analysis showed a significant negative enrichment in NF-kB pathway signaling, while STAT signaling remained largely unaffected (Figure 6D and Supplemental Figure 5). Further analysis revealed that loss of *Uhrf1* leads to decreased levels of p100/p52 (NF-kB1) and p105/p50 (NF-kB2), whereas RelA and RelB protein levels remained largely unchanged (Figure 6E). These finding suggest that Uhrf1 promotes chemokine expression through NF-kB signaling, thereby enhancing immune cell infiltration and sustaining a pro-tumor microenvironment (Figure 6F).

## Discussion

Our study identifies Uhrf1 as a central epigenetic regulator of retinoblastoma progression, demonstrating that its loss prevents tumor formation despite persistent early oncogenic events. These findings build on our understanding of the critical dependency on epigenetic modifications for tumor progression and tumor maintenance following Rb1 inactivation. Through integrative transcriptomic and epigenomic analyses, we establish that Uhrf1 sustains oncogenic transcriptional programs, remodels chromatin accessibility, enforces DNA methylation patterns, and shapes the tumor immune microenvironment. These findings provide new mechanistic insight into how epigenetic dysregulation facilitates retinoblastoma progression and suggest that Uhrf1 inhibition may serve as a promising therapeutic strategy.

A fundamental question in retinoblastoma biology is whether epigenetic dysregulation contributes to tumor initiation, progression, or both. Our findings indicate that Uhrf1 is dispensable for tumor initiation but critical for tumor maintenance. In *Rb1/Rbl1/Uhrf1* TKO retinae, we observed ectopic proliferation at P12, a characteristic of early tumor initiation. However, proliferating cells were absent by P21, contrasting with the persistent proliferation observed in *Rb1/Rbl1* DKO mice. This suggests that tumor initiation can occur independently of Uhrf1, but its loss prevents the transition from dysregulated proliferation to a fully malignant state. This finding has significant implications for our understanding of retinoblastoma progression. While *RB1* inactivation is an early oncogenic event, it is not sufficient to drive tumorigenesis. Instead, cooperating epigenetic mechanisms must sustain aberrant proliferation and prevent differentiation (Zhang et al. 2012). Uhrf1 appears to function at this critical juncture, supporting the hypothesis that Uhrf1 is a major driver of the epigenetic modifications that provide the necessary permissive landscape for tumor progression (Benavente et al. 2014).

An intriguing question is how UHRF1’s epigenetic functions interact with other chromatin-modifying factors. Given that RB1 loss is associated with changes in chromatin accessibility and transcriptional deregulation, it is possible that Uhrf1 cooperates with additional chromatin remodelers to establish a tumor-permissive epigenetic landscape. We have previously reported that the chromatin modifier Hells significantly contributes to retinal tumorigenesis (Zocchi et al. 2020a; Benavente et al. 2014). Here, we found that Uhrf1 loss results in Hells downregulation (Figure 4B). Future studies will be needed to identify potential Uhrf1-interacting partners and determine whether targeting these interactions could provide additional therapeutic benefit.

Beyond its role in epigenetic regulation, we build on the function of Uhrf1 in modulating the tumor immune microenvironment. A recent study uncovered that UHRF1 may downregulate MHC-I, suppressing antigen presentation (Tan et al. 2024). In retinoblastoma, microglia have been implicated in progression by promoting tumor growth, immune suppression, and angiogenesis. Tumor-associated microglia secrete factors that support tumor survival and invasion while simultaneously suppressing anti-tumor immune responses. Studies have shown that retinoblastoma microglia exhibit an enhanced innate response, which may contribute to a more immunosuppressive tumor niche (Xu et al. 2024). Additionally, single-cell transcriptomic analysis of retinoblastoma tumors has identified microglia as key players in shaping the tumor microenvironment, with chemokine signaling being a major driver of their recruitment (Wu et al. 2022a). Importantly, we found that Uhrf1 regulates chemokine secretion via NF-κB signaling to facilitate immune cell recruitment, including microglia, and establish a tumor-supportive microenvironment.

The discovery that UHRF1 loss prevents tumor formation suggests that it may be a promising therapeutic target in retinoblastoma. Unlike broad chromatin-modifying agents, targeting UHRF1 could provide selective tumor inhibition with minimal effects on normal retinal development, as *Rb1/Rbl1/Uhrf1* TKO mice exhibited no significant retinal defects. Additionally, Uhrf1 inhibition not only disrupts tumor cell proliferation but also reprograms the tumor immune microenvironment, reducing microglial infiltration and chemokine signaling. Given the increasing focus on targeting the tumor microenvironment as a cancer therapy, these findings suggest that UHRF1 inhibitors may have dual therapeutic benefits: impairing tumor growth while disrupting tumor-promoting inflammation. Future studies will need to explore small-molecule inhibitors of UHRF1 and assess their potential as a novel treatment strategy for retinoblastoma.

## Materials and methods

### Mouse Models

The *Rb^lox/lox^* mice were obtained from the Mouse Models of Human Cancer Consortium at the National Cancer Institute; the *Rbl1^-/-^* mice were obtained from Dr. Tyler Jacks (Massachusetts Institute of Technology); *Chx10-Cre* mice were obtained from Dr. Connie Cepko (Harvard Medical School); *Z/EG* mice were obtained from Dr. Itsuki Ajioka (Tokyo Medical and Dental University)(Novak et al. 2000). *Uhrf1^lox/lox^* mice were obtained from the European Mouse Mutant Archive, backcrossed to Flp mice for removal of the neo-cassette (tm1c conversion) and then backcrossed to C57BL/6N mice for final Flp removal. *Chx10-Cre Rb^lox/lox^ Rbl1^-/-^* (*Z/EG* negative; *Rb/Rbl1* DKO) mice were monitored weekly for signs of retinoblastoma and anterior chamber invasion for 1 year from the time of birth. Moribund status was defined as the point when tumor cells invaded the anterior chamber and intraocular pressure increased to the point of imminent ocular rupture. Mouse colonies were maintained to include littermate controls carriers and non-carriers of *Chx10-Cre* and/or *Z/EG*. The University of California Irvine Institutional Animal Care and Use Committee approved all animal procedures. Survival curves were generated using GraphPad Prism. Mantel-Cox test was used for statistical analyzes of the curves. Male and female mice were utilized and analyzed in all in vivo studies, with no sex differences identified.

### Real-time RT-qPCR

RNA was isolated from retinal tissue by homogenizing samples using Trizol Reagent and then isolated using chloroform. We used 1 µg of RNA to make cDNA according to SuperScript™III First-strand synthesis system (Invitrogen) manufacturer’s protocol at a reaction volume of 20μl. Quantitative PCR amplification was performed using 1µl of reverse-transcribed product in Power SYBR Green PCR Master Mix (4367659, Life Technologies). Reaction was carried out using 7500 Real-Time PCR system (Applied Biosciences). Data was normalized to endogenous 18S and GAPDH controls and analyzed using the ΔΔ*C*_t_ method. Primers used are listed in Supplemental Table 1.

### Chromatin Immunoprecipitation (ChIP assay)

ChIP assays were performed on Weri retinoblastoma human cell line or postnatal day 0 retinae collected from wild type mice as previously described (Benavente et al. 2013). The antibodies used for chromatin pull-down were anti-E2F1 (3742; Cell Signaling) and rabbit IgG (sc-2027, Santa Cruz Biotechnologies). ChIP DNA was analyzed by qPCR with SYBR Green (Bio-Rad) in ABI-7500 (Applied Biosystems). Primers used are listed in Supplemental Table 1. ChIP experiments were run in triplicate.

### RNA Sequencing

Total RNA was isolated using the RNA Micro Kit (Qiagen). Subsequently, 500 ng of total RNA was used to create the RNA-seq library following the manufacturer’s protocol from purification, mRNA fragmentation through the adenylation of end-repaired cDNA fragments and cleanup (TruSeq Stranded mRNA, Illumina). The collected sample was cleaned with AMPure XP beads (Beckman Coulter) and eluted in 20 μl of 10 mM Tris buffer, pH 8, 0.1% Tween 20. A paired-end 100-bp sequencing run was performed on NovaSeq6000 yielding 348M PE reads with a final library concentration of 2 nM as determined by qPCR (KAPA Biosystem). Statistical analysis for differential gene expression was performed using DESeq2.

### Chromatin profiling

Approximately 50,000 cells were harvested for ATAC-seq for each replicate. Briefly, retinae were dissociated, flow sorted into EGFP- or EGFP+, assessed for cell viability, counted, and washed with PBS. ATAC-seq was performed as previously described ATAC-seq libraries were sequenced with the Illumina NovaSeq6000 using 100bp paired end single indexed run. Raw reads were first QCed (FASTQC) and quality and adapter trimmed using Trimmomatic. Trimmed reads were then aligned to mm10 build of the mouse genome using Bowtie2 (v2.2.5) with alignment parameters: bowtie2 -X 2000– local–dovetail. Potential PCR duplicate reads were removed using *MarkDuplicates* from the Picard tools. Peaks were called in each sample using *MACS2* and further filtered using the ENCODE consensus blacklist regions (http://mitra.stanford.edu/kundaje/akundaje/release/blacklists/mm10-mouse/). Differential peaks were identified using R package *diffbind* across the consolidated peak sets and metrics such as adjusted p values were reported. Heatmap plot was generated using deepTools3.

### Western Blotting

Retinas were homogenized by pellet pestle in RIPA buffer (50mM Tris-HCl, pH=8, 150mM NaCl, 1% NP-40, 0.5 % Sodium deoxycholate, 0.1% SDS, 1mM EDTA) with added protease inhibitor (Mini cOmplete™, Roche). Samples were placed to lyse on ice for 30 minutes and then centrifuged at 14000 RPM for 30 minutes at 4°C. Protein concentration was measured using the BCA protein assay (Pierce™ BCA Protein Assay Kit). 30 µg of total protein was run on SDS-PAGE gels (Mini-PROTEAN, Bio-rad). Gels were transferred onto PVDF membrane (Immobilon-P Membrane, EMD Millipore) using semi-dry transfer apparatus (Bio-rad). Following transfer, membranes were incubated in 3% non-fat dry milk in Tris-Buffered Saline (TBS) with 0.25% Tween (TBS-T) at room temperature for 1 hour. Primary antibodies were diluted in 0.5% non-fat dry milk in TBS-T as follows: 1:1000 anti-UHRF1 (Santa Cruz Biotechnology), 1:4000 anti-beta-Actin (A1978, Sigma-Aldrich). 1:1000 anti-NF-kB1 p105/p50 (Cell Signaling 13586), 1:1000 anti-NF-kB2 p100/p52 (Cell Signaling 4882), 1:1000 anti-RelA/p65 (Cell Signaling 8242), 1:1000 anti-RelB (Cell Signaling 10544). Membranes were incubated in primary antibody overnight at 4°C. The membranes were washed 3 times with TBS-T for 5 minutes per wash then incubated with corresponding secondary antibody for 30 minutes at room temperature. The secondary antibodies were prepared by diluting in 0.5% non-fat dry milk in TBS-T as follows: 1:1000 peroxidase labeled anti-mouse IgG (Vector Laboratories), 1:1000 peroxidase labeled anti-rabbit IgG (Vector Laboratories). After incubation for 30 minutes with secondary antibodies, the membranes were again rinsed 3 times with TBS-T for 5 minutes per wash. Bands were visualized using chemiluminescence (SuperSignal™ West Pico Chemiluminescent Substrate by Thermo Scientific). Band intensity assessed using Image J software.

### Immunohistochemistry

Retinas were isolated in phosphate buffered saline (PBS) and fixed overnight in 4% (w/v) paraformaldehyde. Whole retinas were embedded in 4% (w/v) agarose in PBS. The retinas were then cut into 50 μm slices using a vibratome. Retinal sections were blocked in 5% (v/v) normal donkey, goat or rabbit serum, 0.5% Triton X-100 in PBS for 4 hours at room temperature and then placed in primary antibody in the same blocking solution at 4° overnight. Mouse anti-calbindin antibody (C9848, Sigma) was used at 1:100, rabbit anti-recoverin antibody (AB5585, Millipore) was used at 1:5000 dilution, mouse anti-pH3 antibody (H6409, Sigma) was used at 1:200 dilution, mouse anti-syntaxin antibody (S0664, Sigma) was used at 1:500 dilution, mouse anti-glutamine synthetase (610518, BD Biosciences) was used at 1:100 dilution, rabbit anti-activated caspase 3 antibody (559565, BD Biosciences) was used at 1:1000 dilution, rabbit anti-cone arrestin antibody (AB15282, Millipore) was used at 1:5000 dilution, and sheep anti-chx10 antibody (X1180P, Exalpha Biological) was used at 1:200 dilution. The retinal sections were washed 3 times with PBS and incubated in corresponding secondary antibody diluted 1:500 in respective blocking buffer (goat, rabbit, or donkey) for 1 hour at room temperature in the dark. Again, retina were washed 3 times with PBS and then incubated in 300 μL of Vectastain ABC kit (Vector Laboratories) for 30 minutes at room temperature. Retina washed 3 times in PBS and then placed in 300 μL tyramide Cy3 1:125 in amplification buffer (Perkin Elmer) for 10 minutes. Retinal sections were washed with PBS 3 times and DAPI was added (1:1000 dilution in PBS) for 10 minutes to stain the nuclei. Retina washed 2 times with PBS and mounted on slides. Imaging done using Zeiss Confocal microscope.

### Immunocytochemistry of Dissociated Retina

Retinae were isolated and dissociated using trypsin 100X stock and incubation at 37°C for 5-10 minutes. Trypsin inhibitor and 1000X DNase I was added to sample and incubated at 37°C for 5 minutes. Complete culture medium was then added, and cells were transferred to chamber slides. The chamber slides were prepared with 1X poly-L-lysine incubated for 5 minutes. Cells were allowed to incubate on the slides for 30 minutes at 37°C. Then, media was aspirated, and cells were fixed using 4% (w/v) paraformaldehyde overnight at 4°C. Slides washed with PBS twice. Slides were then incubated with primary antibody overnight at 4°C. Mouse anti-calbindin antibody (C-9848, Sigma) was used at 1:100, rabbit anti-recoverin antibody (AB5585, Millipore) was used at 1:5000 dilution, mouse anti-pH3 antibody (H6409, Sigma) was used at 1:200 dilution, mouse anti-syntaxin antibody (S0664, Sigma) was used at 1:500 dilution, mouse anti-glutamine synthetase (610518, BD Biosciences) was used at 1:100 dilution, rabbit anti-activated caspase 3 antibody (559565, BD Biosciences) was used at 1:1000 dilution, rabbit anti-cone arrestin antibody (AB15282, Millipore) was used at 1:5000 dilution, and sheep anti-chx10 antibody (X1180P, Exalpha Biological) was used at 1:200 dilution. Slides washed 3 times in PBS then incubated with secondary antibody in 1:500 dilution with corresponding blocking buffer for 30 minutes at room temperature in the dark. Slides were washed 3 times with PBS and incubated in 300 μL of Vectastain ABC kit (Vector Laboratories) for 30 minutes at room temperature. Slides were washed 3 times in PBS and incubated with tyramide Cy3 1:150 in amplification buffer (Perkin Elmer) for 10 minutes. After washing with PBS 3 times, DAPI was added (1:1000 dilution in PBS) for 5 minutes to stain the nuclei. Slides were washed twice with PBS and mounted using gelvatol containing DABCO. Imaging was completed using Life Technologies EVOS microscope.

### Electroretinography (ERG)

For scotopic ERG measurement, mice were dark adapted for 24 h prior to ERG recording. Each group consisted of four 5-week-old mice. Under dim red light, mice were anesthetized by intraperitoneal injection of a cocktail consisting of 20 mg/ml ketamine and 5 mg/ml xylazine in phosphate-buffered saline at a dose of 0.1 ml per 20g body weight. Pupils were dilated with 1% tropicamide (Henry Schein, Melville, NY) and applied with 2.5% hypromellose (Akorn, Lake Forest, IL) to keep corneas hydrated.

Contact electrodes were placed on corneas while the reference electrode needle was positioned subdermally between the ears. The a-wave and b-wave responses were recorded followed by a white light stimulus of different flash intensities (−3.3 to 1.7 log cd•s/m2). For each intensity, 3 to 20 recordings were made with the resting intervals for recovery from photobleaching and were averaged for the final amplitude. All ERGs were recorded with the Celeris ophthalmic electrophysiology system (Diagnosys LLC, Lowell, MA) and analyzed with Espion V6 software (Diagnosys LLC). Statistical analysis was performed with paired t-test. Data are represented as means ± SD. The investigator performing ERG measurements was blinded to the mouse genotype.

### Reduced Representation Bisulfite Sequencing (RRBS)

Genomic DNA was extracted from P21 retina of Cre-positive *Rb/Rbl1/Uhrf1* TKO, Cre-negative *Rb/Rbl1* DKO, and Cre-positive *Rb/Rbl1* DKO and eluted in 25 μl of water (Wizard SV genomic DNA purification system A2361 Promega). Samples were processed and analyzed by Zymo Research Epigenetics Services – Methylation Platforms.

### Immune panel

Tumors, from mice never exposed to luciferin, were excised from the eye in sterile conditions, rinsed in HBSS (Hanks’ Balanced Salt Solution, Gibco, Cat# 14025076) and weighted. One spleen was collected for calibration proposes. Tumors were digested in 1mg/mL Collagenase D (11088866001, Millipore Sigma) and 1mg/mL DNase I (79254, Qiagen) in HBSS at 37°C for 10-15 min with gentle agitation. The enzymatic reaction was stopped by adding RPMI (21870-092, Gibco) growth media supplemented with 10% FBS (A5670701, Gibco), and the cell suspension was passed through a 70 µm cell strainer (22363548, Fisher Scientific). Samples were centrifuge at 300 x g for 4 min at 4°C. Cells were resuspended in FACS buffer (PBS + 2% FBS + 2 mM EDTA) and transfer to a 96-well plate. The spleen was minced into small fragments by mechanical force using sterile scalpers and wash with PBS before the cell suspension was passed through a 70 µm cell strainer. Cells were centrifuge at 400 x g for 7 min, and the pellet was resuspended in FACS buffer and transferred to a 96-well plate. Spleen and tumor cells located in the 96-well plate were washed once with FACS buffer. Cells were incubated with Fc block (anti-mouse CD16/CD32, 101302, Biolegend, 1:50) for 10 min at 4°C in the dark to reduce nonspecific antibody binding. Cells were then stained with fluorophore-conjugated antibodies targeting surface markers: 1:50 CD4-PE-Cy7 (100421, Biolegend), 1:50 CD8-PB (100725, Biolegend), 1:50 CD3-PerCP-Cy5.5 (100217, Biolegend), CD69-FITC (104506, Biolegend), 1:100 CD45-PE (157604, Biolegend), 1:45 CD11b-BV650 (101259, Biolegend), 1:50 CD11c-BV785 (117336, Biolegend), and 1:50 LIVE/DEAD staining-AmCyan (L34966A, Invitrogen) for 20 min at 4°C in the dark. After washing twice with FACS buffer, cells were fixed in 0.5% formaldehyde.

Samples were acquired on a BD Fortessa X20 flow cytometer (BD Biosciences), and data were analyzed using FlowJo software (v10.10, BD Biosciences). Compensation was performed using single-stained controls, and gating strategies were defined based on fluorescence-minus-one (FMO) controls and isotype controls.

### Cell culture

The UCI-Rb-1 mouse cell line is derived from a *Rb/Rbl1* DKO retinoblastoma and cultured in Roswell Park Memorial Institute (RPMI) 1640 medium (21870-092, Gibco) supplemented with 1% GlutaMAX (35050061, Life Technologies), 10% fetal bovine serum (FBS; A5670701, Gibco), and 1% penicillin/streptomycin (15140-122, Gibco). UCI-Rb-1 cells were maintained in a humidified incubator at 37°C with 5% CO2 and regularly screened for mycoplasma using a mycoplasma detection kit (13100-01, SouthernBiotech).

### CRISPR KO clones

*Uhrf1* gRNA (gRNA1 sequence: TGATTGAGCTCCCTAAAGAG and gRNA2 sequence: GGTCATGGCCAACTATAACG) was cloned into the doxycycline-inducible CRISPR/Cas9 plasmid TLCV2 (87360, Addgene). pLenti PGK V5-LUC Neo (21471, Addgene) was used for bioluminescence imaging. All lentiviruses were produced by calcium phosphate transfection using HEK293T cells and the lentiviral TLCV2 transfer vector, lentiviral envelope plasmid pCMV-VSV-G (8454, Addgene), and lentiviral packaging plasmid pCMV-dr8.2-dvpr (8455, Addgene). Supernatants containing lentiviral particles were collected 24 and 48 h post-transfection and centrifuged at 1600 g for 10 min at 4°C to remove cell debris. Supernatants were filtered through 0.45 μm PES; filter (ge-0504, Nalgene) and concentrated by ultracentrifugation at 23000 rpm for 1.5 hours at 4°C. Lentiviral pellets were resuspended in cold PBS and stored in aliquots at −80°C. Following lentiviral titering, transduction of target cells was achieved by exposing cells to viral particles in serum-free condition for 6 h using a 0.2 MOI. Selection was carried using either puromycin or G418, depending on the vector. Following selection, *Uhrf1* KO cells were cloned by seeding 0.5 cells per well in a 96-well plate. Single clones were screened for UHRF1 protein levels using Western blotting. Clones with the lowest protein expression UHRF1 levels were selected for in vitro and in vivo studies. KO clones were assessed for CRISPR genome editing using Tracking of Indels by Decomposition (TIDE).

### Chemokine Array

Conditioned media from VC and *Uhrf1* KO UCI-Rb-1 cells was used to profile secreted chemokines using the Proteome Profiler Mouse Chemokine Array Kit (ARY020, R&D Systems) membrane-based immunoassay, following manufacturer instructions.

## Acknowledgements

We thank Kristy Huynh, Kyle Tran, Uyen Tran, Ashley Hadweh, Meagan Khuu, and Yasamin Fazeli for help with mouse genotyping.

## Authors contribution

C.A.B conceived the project. Y.G., M.S-P., L.Z., P.S., S.C.W., S.C., and C.A.B. performed the experiments. J.W. and C.C.C. performed the ATAC-Seq and RNA-seq data analysis. S.S. performed the ERG experiments and ERG data analysis. R.T. guided the immune panel analysis. Y.G., M.S-P, and C.A.B. wrote the manuscript.

